# Beyond queen number: two supergenes coordinate dispersal, mating, and colony founding in the ant *Formica cinerea*

**DOI:** 10.64898/2026.07.28.740873

**Authors:** Giulia Scarparo, Alan Brelsford, Jessica Purcell

## Abstract

Reproductive success often depends on coordinated combinations of morphology, dispersal ability, and mating behavior. Supergenes, genomic regions of suppressed recombination, allow such combinations to be inherited by offspring as a single unit. In ants, independently evolved supergenes control colony queen number, yet few studies have investigated their joint influence on morphology, mating, and colony-founding in sexuals. *Formica cinerea* provides a unique opportunity to address this question because it harbors two supergenes that together produce three queen and male morphs: large monogyne, large polygyne, and small polygyne. Here we show that these supergenes jointly shape an integrated suite of traits across the reproductive cycle. Wing area was primarily associated with the chromosome 3 supergene, with monogyne individuals having larger wings than polygyne individuals. Thorax volume was associated with the chromosome 9 supergene, with small polygyne individuals having reduced thorax volume regardless of social origin. Mating was assortative for both supergenes in large morphs but random in small polygyne queens. Independent colony founding was almost exclusively performed by large monogyne queens; initial egg production was unaffected by mate genotype. These findings show that the two supergenes jointly coordinate dispersal morphology, mate choice, and colony-founding into coherent reproductive strategies, preventing maladaptive intermediate phenotypes.

## Introduction

Reproductive success often emerges from coordinated combinations of traits, including morphology, mating behavior, and life-history strategy; the combinations must be reliably co-inherited to be advantageous. However, if independent alleles control different traits, recombination should generate intermediate phenotypic combinations. Such intermediate phenotypes typically experience reduced fitness and are counterselected. Supergenes, chromosomal regions of suppressed recombination, offer a mechanistic solution to such fitness costs by physically linking the loci controlling different traits, and preventing recombination from breaking apart the coadapted allele combinations that define each reproductive strategy.

The number of known supergenes has grown exponentially in recent years, with many examples linked to alternative life history strategies. In the ruff (*Calidris pugnax*), a supergene controls a suite of differences among male morphs, including plumage, body size, testes mass, and mating behavior (Küpper et al. 2016). In the white-throated sparrow (*Zonotrichia albicollis*), a chromosomal inversion underlies differences in coloration, aggression, and parental care (Tuttle et al. 2016). In *Heliconius* butterflies, supergenes coordinate wing pattern mimicry and disassortative mating preferences (Chouteau et al. 2017).

In ants, independently evolved supergenes control colony social organization across multiple lineages (Kay et al. 2022, Chapuisat 2023, Purcell & Brelsford 2025). In these systems, alternative supergene haplotypes are associated with whether colonies harbor a single reproductive queen (monogyne) or multiple queens (polygyne). Such alternative colony organizations have wide-ranging consequences for a colony’s life cycle. Yet, ant supergene research has remained largely anchored to one focal trait, colony queen number. Where additional phenotypic associations have been documented, they have typically been limited to queen morph (Sigeman et al. 2025; Mona et al. 2025, Darras et al. 2026) and more rarely, to nest founding strategy (Errbii et al. 2024). Whether the broader suite of traits that differ between monogyne and polygyne strategies, including dispersal capacity, mating behavior, and reproductive investment, is functionally integrated and maintained by these supergenes has rarely been tested, except in the Alpine silver ant *Formica selysi* and the red imported fire ant *Solenopsis invicta* (Kay et al. 2022, Chapuisat 2023).

The existence of coordinated phenotypic differences between monogyne and polygyne ant queens was recognized well before supergenes were discovered and is often referred to as polygyny syndrome (Keller 1993). Classical studies described how monogyne and polygyne queens differ in body size, fat reserves, flight muscle, dispersal behavior, and are often associated indirectly with alternative colony-founding strategies. Large monogyne queens are assumed to found colonies independently, and may undertake long-distance dispersal flights. They then metabolize fat reserves and flight muscles to provision their first brood in the absence of workers. Smaller polygyne queens, by contrast, are assumed to disperse locally and join or co-found colonies near their natal nest. These observations suggested that alternative social strategies involve integrated life-history syndromes. However, this earlier work was largely descriptive and anecdotal and predated the discovery of supergenes (for instance, most evidence on *S. invicta* life history was accumulated before the supergene was characterized by Wang et al. 2013, but see for example Avril et al. 2019a; De Gasperin et al. 2024 for more recent work directly linking reproductive traits to supergene haplotypes in *F. selysi*). It therefore remains unclear to what extent these trait differences are functionally coordinated at the genomic level and whether the supergenes now known to control social organization also control a broader reproductive phenotype.

*Formica cinerea* provides an ideal system to test whether supergenes integrate multiple reproductive traits. In this species, two supergenes control alternative phenotypes: one on chromosome 3 is associated with colony social organization (monogyne vs. polygyne), and one on chromosome 9 is associated with sexual body size, with three distinct queen and male morphs existing in natural populations (Scarparo et al. 2023). Queens carrying the 9a9a genotype on chromosome 9 are large and defined as macrogynes. The largest macrogynes are monogyne and harbor exclusively M haplotypes (M_A_ and M_D_) on chromosome 3. Polygyne macrogynes are approximately 5% smaller and carry at least one P_1_ haplotype on chromosome 3. Microgynes are up to 20% smaller than macrogynes and are exclusively found in polygyne colonies; they harbor at least one P_2_ haplotype on chromosome 3 and at least one 9r haplotype on chromosome 9 (Scarparo et al. 2023). The strong linkage disequilibrium between P_2_ and 9r is maintained by the high mortality rate of haplotype-mismatched individuals (Scarparo et al. 2026). Males, that are haploid, also exhibit body size variation associated with supergene haplotype: P_1_-9a males are 2.6% larger than M_A_-9a males, while P_2_-9r males are 8.6% smaller than 9a males. M_D_-9a males are extremely rare but expected to be as large as M_A_-9a males (Scarparo et al. 2023). This three-morph system raises a fundamental question that a simple monogyne–polygyne comparison cannot answer: do polygyne macrogynes and microgynes represent divergent reproductive strategies, or do they share the same polygyne life history despite their size difference?

Here, we investigate whether supergenes maintain integrated reproductive phentypes by coordinating multiple traits across the three queen and male morphs. If so, we predict that supergene-associated morphs should exhibit correspondingly different dispersal strategies and mating behaviors reflecting these founding modes.

We test these predictions by examining queens and males carrying different supergene haplotypes for differences in: (1) dispersal ability, measured via wing and thorax morphology; (2) mate choice patterns, assessed through genotype frequencies of field-collected queens and the contents of their spermatheca; (3) queen foundress types (independent vs. dependent founding), inferred from genotypes of queens attempting independent colony establishment; and (4) reproductive investment, examined through egg production. We predict that large monogyne (MM) queens will exhibit superior dispersal potential, while smaller polygyne queens (P_1_ and P_2_ carriers) will show reduced dispersal ability and mate proximally to the maternal nest. We expect dispersal ability to correlate with mating strategy: queens and males that engage in dispersal flights should preferentially mate with similarly dispersing partners, thereby generating assortative mating. Among queens attempting independent colony foundation, we expect to find predominantly MM queens, reflecting the body size requirements for solitary founding. Finally, we predict that mate genotype does not affect the initial egg investment of newly mated queens, based on findings in a closely related species (Avril et al. 2019b, Choppin et al. 2026). Together, these predictions test whether supergenes functionally coordinate dispersal behavior, mate choice, and reproductive dynamics, thereby contributing to the maintenance of the complex social polymorphism in *F. cinerea*.

## Material and Methods

### Sample collection

*Formica cinerea* queens, gynes (unmated queens), and males were collected from the Aosta Valley and Piedmont regions of Italy between late June and early July across multiple collection years (2019-2022). Mated queens were assigned to one of two categories depending on how they were collected: those taken directly from existing colonies along with several workers were referred to as mature queens (n=65), and those caught during mating flights while clinging to vegetation (n=15) or moving across the ground in search of nest sites or under rock without workers (n=63) (referred to as newly mated queens). If newly mated queens were collected under rocks we checked carefully for eggs. Following the collection, each queen was kept individually in a glass tube. Newly mated queens received water only, whereas mature queens were supplied with a sugar-water solution and kept alongside three to four workers from their colony. We checked and counted brood produced by each queen every two days after sampling. Queens were maintained alive for an average of two months before being preserved in 100% ethanol for subsequent genetic analysis.

Colony fragments containing gynes and males were also collected from the same sites during the same years and seasons as the queens. These samples were previously sequenced and described in Scarparo et al. 2023.

### Dispersal abilities

We investigated the dispersal abilities of gynes and males using the indirect morphological proxies of wing area and thorax volume. Wing length (WL) and wing width (WW) were measured on either the right or left forewing (chosen at random) for 331 males and 149 gynes. Prior to measurement, wings were mounted on microscope slides to ensure they lay completely flat. Wing area (WA) was then approximated using the formula for the area of a triangle: WA = ½ × WL × WW.

Thorax length (TL) and thorax width (TW) were measured for 211 males and 100 gynes. Thorax volume (TV) was estimated by treating the thorax as an ellipsoid with height equal to width, using the formula: TV = (4/3) × π × (TW/2)² × (TL/2) (Turlure et al. 2010).

All measurements were taken using a Leica DMC2900 camera mounted on a Leica S8APO stereomicroscope at 10× magnification. Head width measurements for the same individuals, obtained using the same microscope at 25× magnification, were previously published in Scarparo et al. (2023), as were their supergene genotypes, which had been determined via double-digest restriction-site associated DNA sequencing (ddRAD-seq).

### Genotype distribution among newly mated and mature queens and their mates

We collected and genotyped 78 newly mated queens and 65 mature queens. Queens sampled between 2019 and 2021 were analyzed using RAD-seq as part of a previous study (Scarparo et al. 2023), while those collected in 2022 were genotyped using PCR–restriction fragment length polymorphism (PCR-RFLP) assays described in Scarparo et al. 2026. To assess whether queens with specific haplotype combinations preferentially mate with males carrying the same haplotypes, we dissected their spermathecae to determine the genotypes of their mates. In total, we successfully dissected and genotyped the spermathecae of 122 queens. In queens that laid eggs (n=88), mate genotypes were further validated by analyzing egg genotypes. Spermatheca contents and eggs were genotyped via PCR-RFLP. These genotyping data were originally generated to assess the nature of linkage disequilibrium between P_2_ and 9r in Scarparo et al. 2026.

To assess whether male genotype affects queen fecundity, we reared 57 newly mated queens under controlled conditions, recording the number of eggs laid through daily observation. Queens were maintained until the first workers eclosed (approximately four to six weeks), at which point they were euthanized. We defined the initial egg number as the total number of eggs recorded on the last day before larvae were first observed. Male genotype was inferred by dissecting the queen’s spermatheca and genotyping the stored sperm via PCR-RFLP, as described above.

### Statistical analyses

Given the genotypic complexity of the supergene system and the hierarchical dominance relationships among haplotypes (M_A_ and M_D_ < P_1_ < P_2_ for chromosome 3; 9a < 9r for chromosome 9), we combined genotypes to increase statistical power. For chromosome 3, individuals carrying at least one P_2_ haplotype (e.g., P_2_P_2_, P_2_P_1_, MP_2_) were grouped as P_2_; individuals carrying at least one P_1_ haplotype (but not P_2_) were grouped as P_1_ (e.g., P_1_P_1_, MP_1_); and individuals carrying neither P_1_ nor P_2_ were grouped as M, regardless of whether they were M_A_ or M_D_. This grouping was applied consistently to analyses of both morphological traits and mating patterns. For the analysis of morphological traits only, a similar approach was applied to chromosome 9: individuals carrying at least one 9r haplotype (9a9r or 9r9r) were grouped as 9r, and those lacking 9r were grouped as 9a.

To test whether supergene haplotypes on chromosomes 3 and 9 are associated with wing area and thorax volume in males and gynes, we fitted several linear mixed models. Models were fitted independently for each sex (gynes and males) and for each chromosome (3 and 9). In each model, log-transformed wing area or thorax volume was the response variable, with genotype category and log-transformed head width as fixed effects and colony ID as a random effect. Models were fitted using the lmer function from the lme4 package (Bates et al. 2015) in R, and post-hoc pairwise comparisons with Tukey correction were obtained using the emmeans package (Lenth et al. 2019).

To distinguish active assortative mate choice from mating patterns that simply reflect the relative availability of male morphs, we estimated population-level male haplotype frequencies independently of the mating data. Because each male-producing colony produces males with the same supergene haplotypes with very rare exceptions (Scarparo et al. 2023, Palanchon 2026), we estimated relative male morph frequencies by counting the number of colonies producing each male genotype, rather than the number of individual males sampled, to avoid bias from uneven sampling effort across colonies. For chromosome 3, this yielded frequencies of 51.7% M, 24.1% P_1_, and 24.1% P_2_ (45, 21, and 21 colonies, respectively); for chromosome 9, 75.9% 9a and 24.1% 9r. However, this colony-based approach assumes broadly comparable male output across colonies, and does not account for unequal production of males among colonies or years. For each queen genotype class, we tested whether the observed distribution of mate genotypes differed from these population-level male frequencies using a chi-square goodness-of-fit test, with p-values estimated by Monte Carlo simulation (10,000 replicates) to account for small, expected cell counts. To identify which crosses contributed most to significant deviations, we examined the corresponding Pearson residuals. Double-mated queens were excluded from this analysis. We then tested whether double mating with males carrying different genotypes was more common in heterozygous queens than in homozygous queens, as previously observed in *F. selysi* (Avril et al. 2019b), by running a Fisher’s exact test with the following levels: heterozygous/homozygous and single-mated/double-mated, independently for chromosomes 3 and 9.

To test whether newly mated queens collected during mating flights and independent colony founding preferentially carry monogyne-associated genotypes, we performed a binomial test, defining “success” as the number of queens carrying monogyne-associated genotypes, and comparing it against an expected probability of 0.45, reflecting the estimated relative frequency of MM gynes in the population. This estimated frequency was calculated by counting the number of gyne-producing colonies of each genotype (MM, XP1, XP2), following the same approach used for the male morph frequencies above (mating pattern analysis). Following Scarparo et al. (2023), monogyne colonies are expected to produce MM gynes, while polygyne colonies are expected to produce gynes with at least one P haplotype. Finally, to test whether newly mated queens mated to males with different genotypes differ in egg laying, we fitted a GLM with quasipoisson error structure to account for overdispersion, with the initial egg number recorded before the appearance of the first larvae as the response variable and mate genotype as a fixed effect. Since all newly mated queens carried monogyne-associated genotypes, queen genotype was not included in the models. This was tested independently for chromosome 3 and chromosome 9.

## Results

### Dispersal phase

Our wing and thorax measurements for gynes and males revealed a striking pattern, with each trait being primarily driven by a different supergene (full statistical details in Table 1). Across all individuals, head width was strongly and positively correlated with both wing and thorax size (p < 0.0001 in all cases, Figure 1).

**Figure 1.**
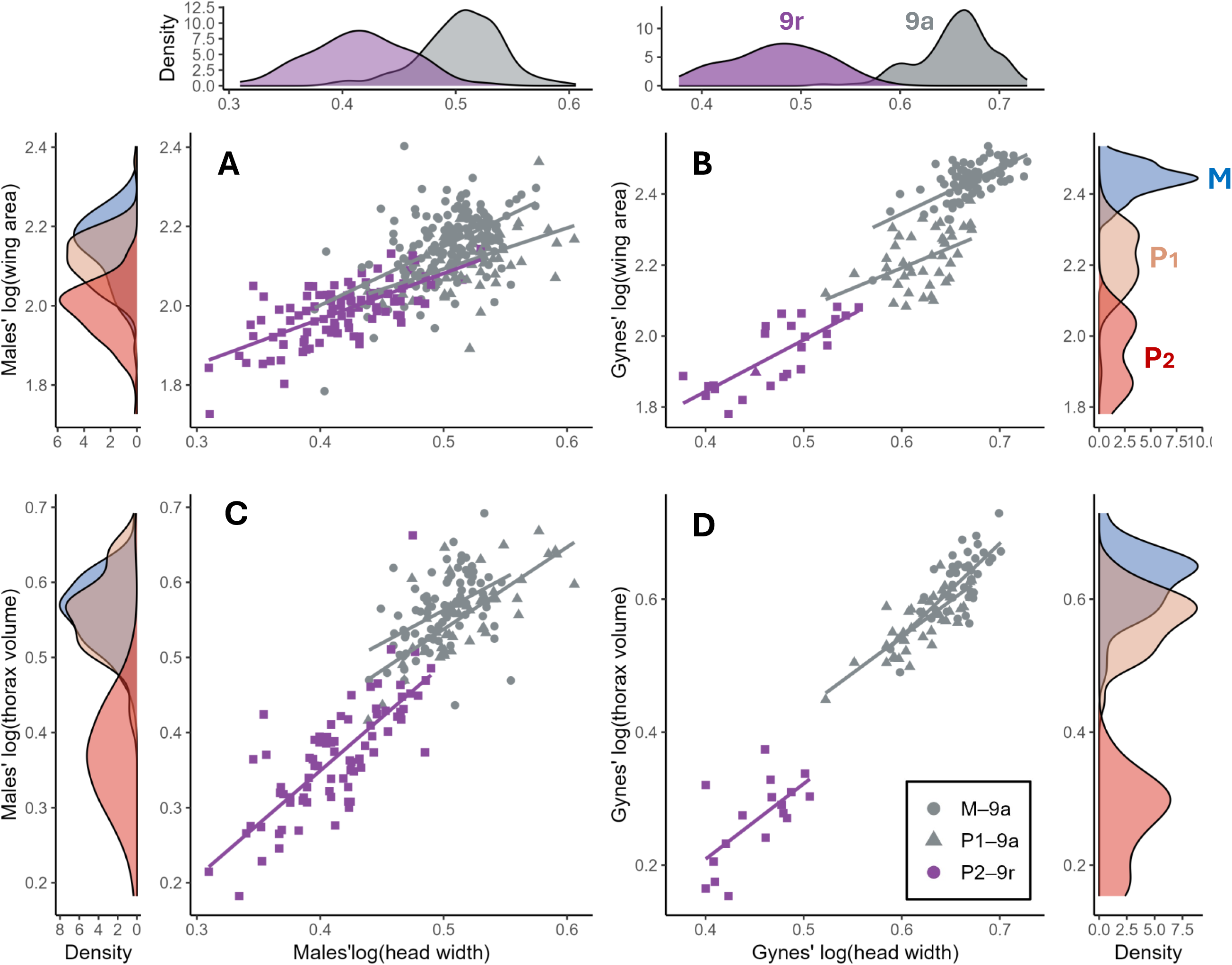
Relationships between body size (log head width) and log wing area (A, B) and log thorax volume dispersal-related morphology (C, D) in males (right column) and gynes (left column) of *Formica cinerea*. Colors indicate chromosome 9 haplotypes (9a in grey, 9r in purple), and shapes indicate chromosome 3 genotypes (M, P_1_, P_2_). Regression lines are shown for each haplotype. Marginal density plots illustrate the distribution of head width (top panels) relative to chromosome 9 and dispersal-related traits (side panels) relative to chromosome 3.

**Table 1.** Pairwise contrasts from linear mixed models for wing area and thorax volume. Estimates are log-scale differences. Significant comparisons (p < 0.05) are shown in bold.

| Sex | Contrast | Estimate | SE | df | t. ratio | p |
| --- | --- | --- | --- | --- | --- | --- |
| Wing area — chromosome 3 |  |  |  |  |  |  |
| Male | M - P<sub>1</sub> | 0.059 | 0.013 | 97.3 | 4.63 | **<0.0001** |
| Male | M - P<sub>2</sub> | 0.074 | 0.015 | 116.1 | 5.12 | **<0.0001** |
| Male | P<sub>1</sub> - P<sub>2</sub> | 0.015 | 0.018 | 127.3 | 0.86 | 0.67 |
| Gyne | M - P<sub>1</sub> | 0.15 | 0.016 | 43.2 | 9.37 | **<0.0001** |
| Gyne | M - P<sub>2</sub> | 0.2 | 0.029 | 126.6 | 6.8 | **<0.0001** |
| Gyne | P<sub>1</sub> - P<sub>2</sub> | 0.048 | 0.025 | 113.1 | 1.95 | 0.13 |
| Wing area — chromosome 9 |  |  |  |  |  |  |
| Male | 9a - 9r | 0.067 | 0.016 | 124 | 4.27 | **<0.0001** |
| Gyne | 9a - 9r | 0.105 | 0.04 | 81.2 | 2.61 | **0.01** |
| Thorax volume — chromosome 3 |  |  |  |  |  |  |
| Male | M - P<sub>1</sub> | 0.028 | 0.011 | 43.7 | 2.65 | **0.03** |
| Male | M - P<sub>2</sub> | 0.095 | 0.012 | 79.7 | 7.76 | **<0.0001** |
| Male | P<sub>1</sub> - P<sub>2</sub> | 0.067 | 0.015 | 82.9 | 4.6 | **<0.0001** |
| Gyne | M - P<sub>1</sub> | 0.018 | 0.012 | 27.7 | 1.55 | 0.285 |
| Gyne | M - P<sub>2</sub> | 0.107 | 0.028 | 78.2 | 3.82 | **0.0008** |
| Gyne | P<sub>1</sub> - P<sub>2</sub> | 0.089 | 0.024 | 68 | 3.65 | **0.0015** |
| Thorax volume — chromosome 9 |  |  |  |  |  |  |
| Male | 9a - 9r | 0.089 | 0.013 | 90.3 | 7.14 | **<0.0001** |
| Gyne | 9a - 9r | 0.085 | 0.025 | 64.9 | 3.45 | **0.001** |

Wing area appears to be primarily associated with the supergene on chromosome 3, as supported by the pattern of pairwise differences (Figure 1A-B). In both sexes, M individuals had significantly larger wings than both P_1_ and P_2_ individuals (males p < 0.0001; gynes: p < 0.0001), while P_1_ and P_2_ individuals did not differ significantly from each other despite carrying different chromosome 9 haplotypes (males: t= 0.86, p = 0.67; gynes: t= 0.13, p = 0.13). Based on model estimates, M males had approximately 6% larger wings than P_1_ males and 7.7% larger wings than P_2_ males (Figure 1A), whereas MM gynes had approximately 16.2% larger wings than gynes carrying at least one P_1_ and 22.1% larger wings than P_2_ gynes (Figure 1B, Table 1). A secondary effect of chromosome 9 on wing area was also detected in both sexes: 9r individuals had significantly smaller wings than 9a9a individuals (males: t = 4.27, p < 0.0001; gynes: t = 2.61, p = 0.01), likely reflecting the linkage disequilibrium between P_2_ and 9r rather than an independent effect of chromosome.

Thorax volume is primarily associated with the supergene on chromosome 9 (Figure 1C-D). In both sexes, individuals carrying at least one 9r haplotype had significantly smaller thoraxes than 9a individuals (males: t = 7.14, p < 0.0001, Figure 1C; gynes: t = 3.45, p = 0.001, Figure 1D), corresponding to a reduction of approximately 9% in both males and gynes based on model estimates. Results for chromosome 3 mirrored this pattern in both sexes, consistent with the tight association between the P_2_ and 9r haplotypes, and are reported in full in Table 1. No difference in thorax volume was detected between MM gynes and those carrying at least one P_1_ haplotype (Table 1).

### Mating phase

Partial assortative mating has previously been detected in *F. selysi* (Avril et al. 2019a) and *S. invicta* (Ross and Keller 1995), and our results in *F. cinerea* show a similar pattern. Analyses of mating patterns revealed significant deviation from random mating with respect to both chromosome 3 and chromosome 9 (p < 0.0001 in both cases) (Figure 2). However, the signal of assortative mating is not uniform across genotypes in *F. cinerea*.

**Figure 2.**
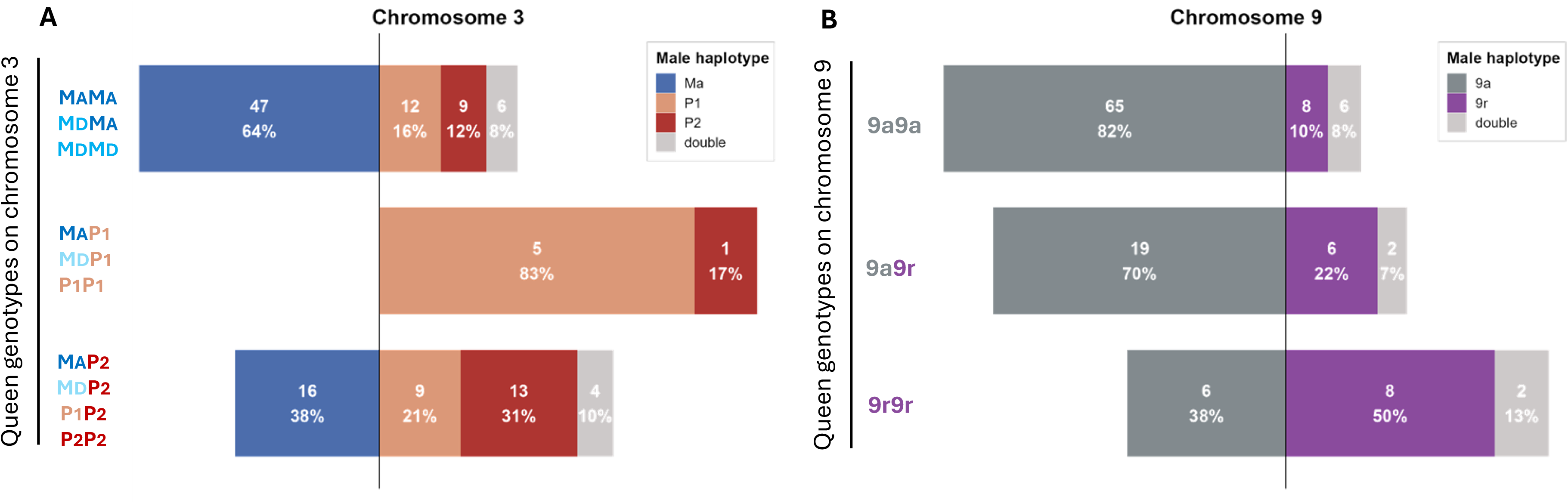
Mating patterns of *F. cinerea* with respect to supergene haplotypes on chromosome 3 (A) and chromosome 9 (B). Bars represent the proportion of male haplotypes mated by each queen genotype class, with raw counts and percentages shown within each bar.

For each queen genotype, we tested whether the distribution of mate genotypes deviated from expectations based on population-level male morph frequencies (chromosome 3: M = 51.7%, P1 = 24.1%, P2 = 24.1%, and chromosome 9: 9a = 75.9%, 9r = 24.1%) using chi-square goodness-of-fit tests with simulation-based p-values”. With respect to chromosome 3, MM queens (p = 0.016) and MP_1_/P_1_P_1_ queens (p = 0.003) showed significant deviations from these expectations, while MP_2_/P_2_P_2_/P_1_P_2_ queens did not (p = 0.32). Standardized residuals indicated that MM queens mated with M males more often than expected (residual = 1.99) and with P_2_ males less often than expected (residual = −1.83), while MP_1_/P_1_P_1_ queens mated with P_1_ males far more often than expected (residual = 2.95) despite M being the most abundant male morph in the population (residual for M = −1.76) (Figure 2A). On chromosome 9, the largest 9a9a (p = 0.009) and the smallest 9r9r queens (p = 0.007) showed mate distributions that deviated significantly from the population-level male frequencies, whereas 9a9r heterozygous queens did not (p = 1). Standardized residuals showed that 9a9a queens mated with 9a males more often than expected (residual = 1.29) and with 9r males less often than expected (residual = −2.29), while 9r9r queens showed the opposite pattern, mating with 9r males far more often than expected (residual = 2.51) despite 9a being the more abundant male morph (residual for 9a = −1.42). After accounting for the estimated frequencies of 9a and 9r males available in the population, queens carrying the 9a9r genotype mate with each male genotype proportionally, consistent with random mating rather than assortative mating (Figure 2B). These patterns indicate assortative mating rather than mating proportional to male morph availability, consistent with the haplotype-specific preferences described above.

Multiple mating was generally moderate in our sample, with at least 10 out of 122 queens (8.2%) mated with two (or more) males. This estimate is likely conservative, as our PCR-RFLP assays of spermatheca contents do not distinguish cases where a queen has mated multiple times with males carrying the same haplotype. Contrary to what has been observed in *F. selysi*, our data show no evidence that queens heterozygous on chromosome 3 or chromosome 9 mate multiply more often than their homozygous counterparts (chromosome 3: p = 0.75; chromosome 9: p = 0.68).

### Colony foundation phase

Of the 78 newly mated queens collected, 75 carried MM-9a9a genotypes exclusively; of the three remaining queens, two were P_2_P_2_-9r9r and one P_1_P_1_-9a9a. In contrast, all but three of the mature queens collected carried at least one P haplotype. However, the sample of mature queens is not representative of population-level queen genotype frequencies. To minimize disturbance, we deliberately restricted sampling to colonies containing multiple queens and consistently collected fewer individual queens than observed within each colony; the only exceptions were three mature MM queens collected from monogyne colonies. Although genotype distributions between newly mated and mature queens are therefore not directly comparable, some patterns emerge. The proportion of newly mated queens carrying monogyne-associated genotypes and attempting independent colony foundation was 96% (75/78), significantly higher than expected based on the estimated relative frequency of 45% MM gynes in the population (binomial test: p < 0.0001) (Figure 3). In contrast, observations from mature queens are consistent with previous genomic evidence (Scarparo et al. 2023), showing that P haplotypes (P_1_ and P_2_) are associated with multi-queen colonies (Figure 3).

**Figure 3.**
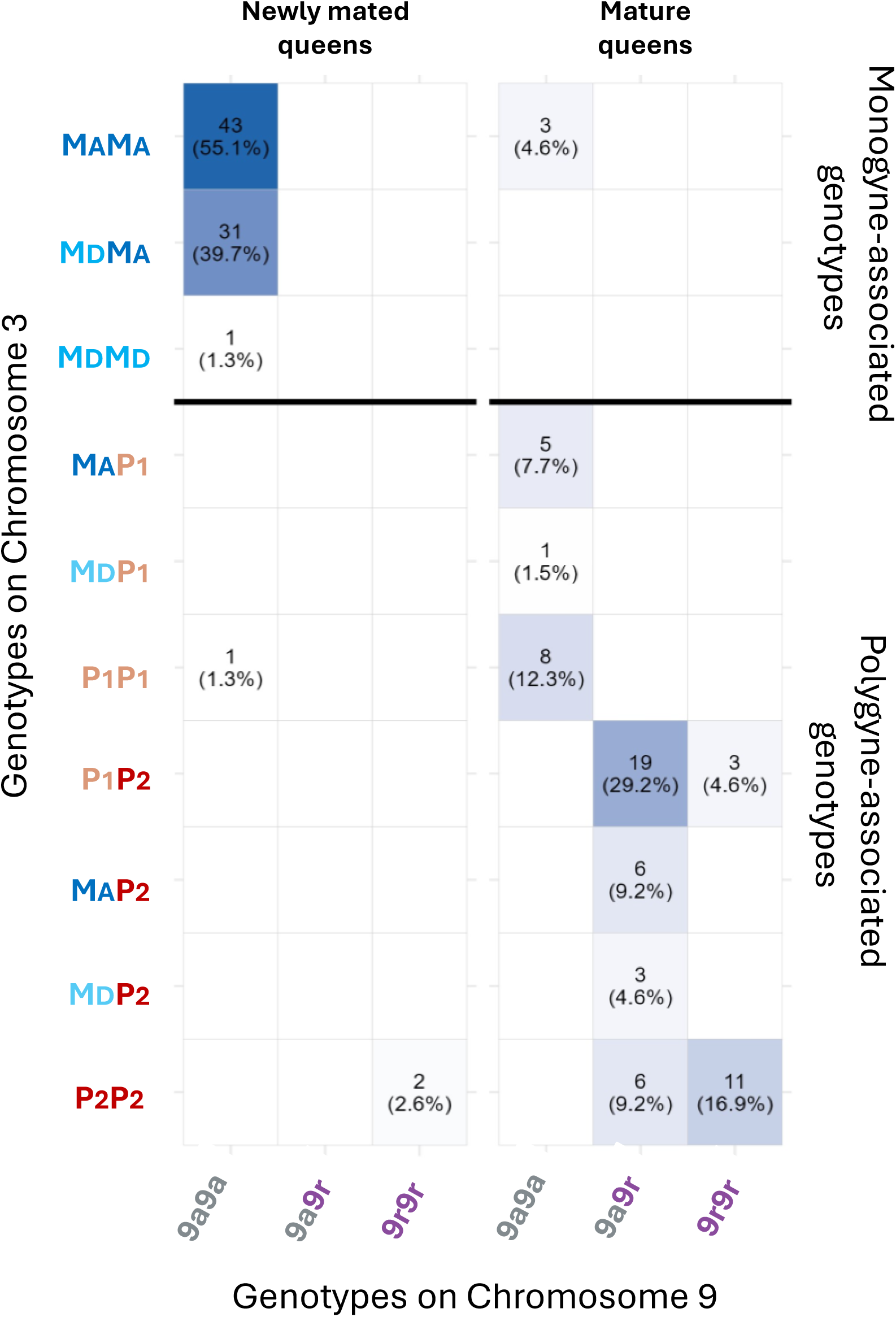
Heatmap of supergene genotype combinations on chromosomes 3 and 9 in newly mated and mature queens. Newly mated queens were sampled during mating flights or colony founding attempts, whereas mature queens were collected from established colonies. Cell color indicates genotype frequency (darker shades represent higher frequencies), and each cell shows the number of queens with the corresponding percentage in parentheses. Blank cells indicate unobserved genotype combinations. The horizontal line separates monogyne-associated genotypes (suggested to be linked to independent colony founding) from polygyne-associated genotypes (suggested to be linked to dependent colony founding).

The number of eggs produced during colony foundation in the lab did not differ significantly among newly mated queens mated with males carrying different haplotypes on chromosome 3 (likelihood ratio χ² = 2.82, df = 2, p = 0.24) or 9 (likelihood ratio χ² = 0.83, df = 1, p = 0.36).

## Discussion

The two supergenes in *Formica cinerea* do not simply control colony queen number or queen body size. Instead, they appear to coordinate an integrated reproductive strategy linking morphology, dispersal, mating patterns, and colony foundation. Our results show that individuals carrying alternative supergene haplotypes differ not only in body size but also in wing area and thorax volume, traits that are closely associated with dispersal ability in flying insects. These morphological differences likely translate into contrasting dispersal strategies that ultimately shape mating and colony foundation dynamics (Figure 4).

**Figure 4.**
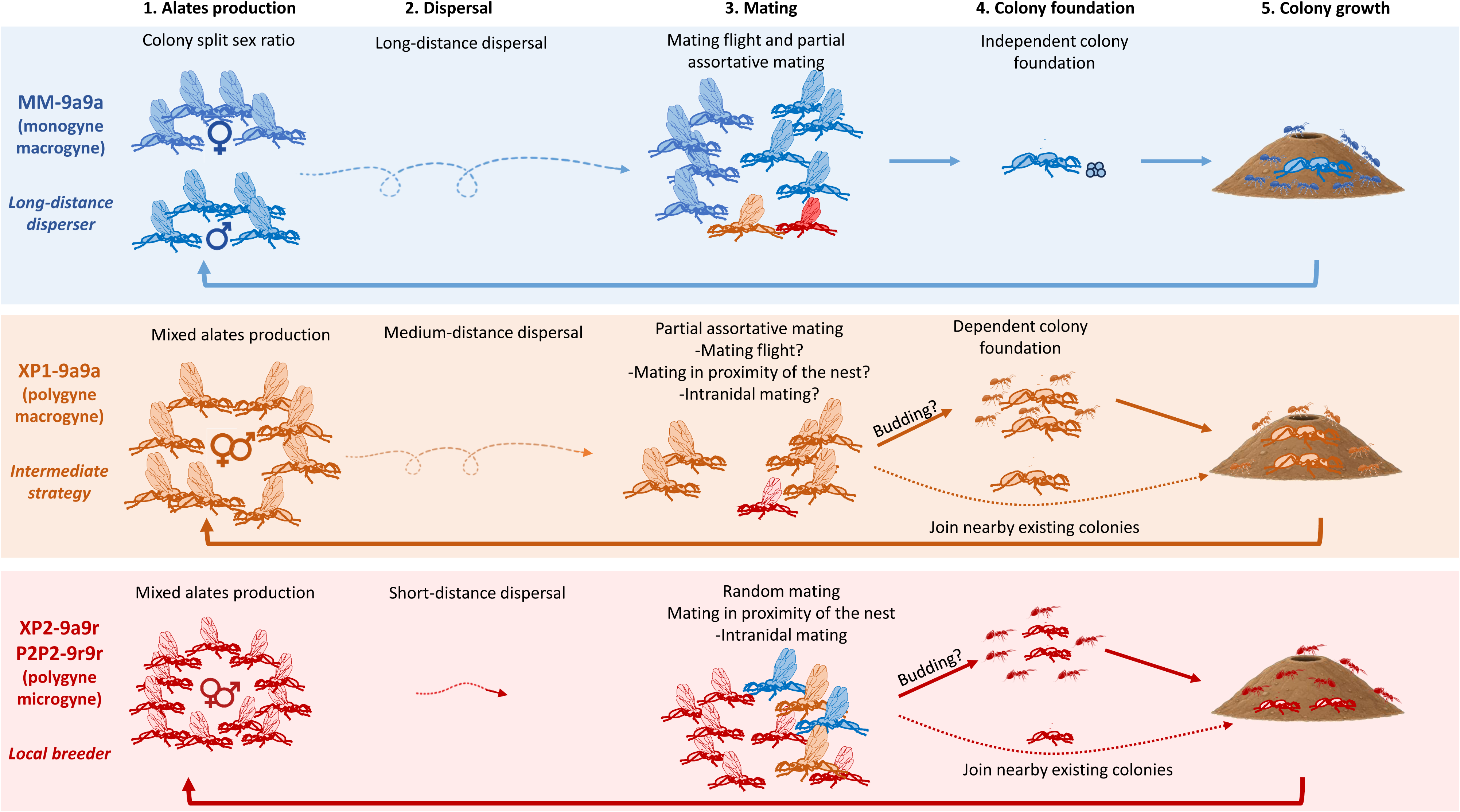
Overview of suggested reproductive strategies associated with supergene haplotypes on chromosomes 3 and 9 in *F. cinerea*.

Two lines of evidence support this interpretation. First, mating patterns are assortative with respect to supergene haplotypes, with large monogyne (MM-9a9a) and large polygyne (XP_1_-9a9a) queens preferentially mating with M-9a and P_1_-9a males, respectively, although queens carrying the P_2_ haplotype or the 9a9r genotype mate randomly. Second, genotype frequencies among newly mated queens attempting to establish a new colony were not random. Together, these results suggest that the supergenes influence reproductive outcomes across multiple stages of the life cycle. In combination with previous work in *Formica selysi* and *Solenopsis invicta* (see Kay et al. 2022, Chapuisat 2023, and references therein), our findings provide some of the strongest evidence to date that supergenes can organize entire reproductive phenotypes in social insects.

### Dispersal phase

In *F. cinerea*, queens and males occur in three morphs: a large monogyne morph (macrogynes and macraners), a large polygyne morph (macrogynes and macraners), and a small polygyne morph (microgynes and micraners). Before this study, however, it remained unclear whether the supergenes associated with social organization also influence morphological traits related to dispersal. Overall, we find that polygyne-associated haplotypes (P_1_ and P_2_) and the 9r haplotype are associated with reduced wing and thorax size. These findings broadly mirror patterns observed in the closely related species *Formica selysi* (De Gasperin et al. 2024). In that system, a single supergene on chromosome 3 (Purcell et al. 2014) controls both social organization and morphological variation, with individuals carrying the P haplotype exhibiting reduced wing and thorax size. However, the more complex supergene system in *F. cinerea* where two supergenes interact generates a broader range of morphologies and sharper differences between morphs. Reduced wing area and thorax volume are commonly associated with limited dispersal ability (Sekar 2012). Individuals with larger wings and thoraxes are typically better suited for long-distance dispersal flights, whereas individuals with smaller wings and thoraxes often disperse over shorter distances and remain close to their natal population (Sundström 1995; Ross and Keller 1995). Based on these patterns, we propose that large monogyne individuals (MM–9a9a gynes and M–9a males) represent the primary dispersers in the population. In contrast, individuals carrying the P_2_–9r combination are likely more philopatric. Individuals carrying P_1_-9a haplotypes show intermediate morphology, and further research is needed to determine which strategies they employ.

Most interestingly, here, we show that the two supergenes affect different components of flight morphology. The supergene on chromosome 3 primarily influences wing area, with individuals carrying P_1_ and P_2_ haplotypes having significantly smaller wings than M males and MM queens. In contrast, the supergene on chromosome 9 has a stronger effect on thorax volume, with 9r individuals having smaller thoraxes than 9a males and 9a9a queens. A recent study (Scarparo et al. 2026) showed that the P_2_ haplotype on chromosome 3 and the 9r haplotype on chromosome 9 assort independently during meiosis, yet exhibit strong linkage disequilibrium in adults. Scarparo et al. (2026) traced this pattern to intrinsic developmental mortality rather than extrinsic selection on adult phenotypes: individuals carrying mismatched combinations (whether between wing and thorax size or between dispersal morphology and social background [e.g., monogyne–small-body combinations]) largely fail to survive development, rather than surviving to eclosion and being selected against through reduced mating or colony-founding success. While we do not yet know the exact developmental mechanism causing the P_2_–9r combination to be overrepresented among adults relative to mismatched combinations, nor whether the few mismatched individuals that survive to adulthood face further reduced fitness, the strong LD between P_2_ and 9r effectively couples otherwise independent supergenes into a single functional unit controlling dispersal and reproductive strategy.

### Mating phase

Differential dispersal abilities are often associated with distinct mating strategies (Sundström 1995; Ross and Keller 1995). Monogyne individuals that are better equipped for long-distance dispersal not only disperse far from the natal nest but are also thought to participate in mating flights, whereas polygyne individuals, which tend to be more philopatric, mate in the vicinity of their natal colony (Fontcuberta et al. 2021). This coupling between social form, dispersal morphology, and mating strategy is expected to produce assortative mating even in the absence of mate choice.

Our results support this prediction, although the strength of assortative mating varies across supergene haplotypes. Monogyne MM queens mate most often with M males and less frequently than expected with P_2_ males (Figure 2A). Similarly, queens carrying at least one P_1_ haplotype, but not P_2_, tend to mate most often with P1 males, though we acknowledge that sample sizes for these crosses remain limited. A parallel signal emerges for the supergene on chromosome 9: the smallest queens (9r9r) mate most often with small 9r males, while large 9a9a queens mate most often with large 9a males, consistent with size-assortative mating driven by differential dispersal.

Comparable patterns have been reported in *F. selysi* (Avril et al. 2019a), where queens of both social forms mate most often with males carrying the corresponding supergene haplotype, with gene flow between social forms detected in approximately 20% of matings (Avril et al. 2019a; Fontcuberta et al. 2021). However, in *F. selysi* the observed assortative mating does not appear to be driven by active mate preference (Avril et al. 2019b). This contrasts with observations in fire ants, where both queen morphs mate most often with males carrying the monogyne-associated haplotype, and with cases in some butterflies (Chouteau et al. 2017) and the white-throated sparrow (Tuttle et al. 2016), where mating is completely disassortative. Assortative mating was, however, not detected in queens carrying the P_2_ haplotype or in heterozygous 9a9r queens. Because most P_2_-carrying queens in our dataset are also 9a9r heterozygotes, these two results likely reflect largely overlapping sets of individuals rather than two independent signals. These 9a9r queens have significantly smaller bodies than 9a9a queens but larger bodies than 9r9r queens (Scarparo et al. 2023), suggesting an intermediate phenotype that may allow them to mate randomly with males of different sizes. Double mating appears to be rare, observed in only 8.2% of queens, a rate comparable to that reported in *F. selysi* (Avril et al. 2019a). Unlike in *F. selysi*, where heterozygous queens are more frequently double-mated, we find no such association in our dataset.

These patterns also offer indirect insight into the mating context. Of the 15 newly mated queens caught during a mating flight, all were MM; of these, 10 were mated to an M male and 4 to a P_2_ male (one failed amplification). This pattern suggests that MM queens do frequently engage in mating flights, while queens carrying a P haplotype do not, although this inference awaits confirmation from larger sample sizes. By exclusion, we suggest that queens carrying a P_2_ haplotype, given their reduced flight capabilities, are likely to remain near the maternal nest and mate locally. Supporting this, we observed that P_2_-9r microgynes and micraners collected from the same colony will occasionally mate when kept together, suggesting that intranidal mating is one of the strategies employed by these smaller individuals (pers. obs.). However, because P_2_ males make up nearly all (or all) males within nests where P_2_ is present, that finding P_2_ gynes mated with M or P_1_ males implies that these queens mated outside their natal nest (Scarparo et al. 2023; Palanchon 2026).

On the male side, P_2_–9r micraner males, perhaps owing to their subtle morphological differences from macraners, appear capable of engaging in mating flights alongside M–9a males. Although none of the 15 newly mated queens caught during a mating flight had mated with P_1_–9a males, our extended dataset, which includes newly mated queens collected outside mating flights and mature queens, shows that P_1_–9a males do occasionally mate with MM queens, suggesting that this male morph can also participate in mating flights. Taken together, our results indicate an asymmetry between the sexes: queens may have evolved more rigid, morph-specific mating strategies, whereas males of all genotypes retain the flexibility to engage in both mating flights and local mating, thereby enabling continuous gene flow between morphs. Similar male-mediated gene flow was also observed in *F. selysi* (Fontcuberta et al. 2021).

### Colony foundation phase

Following mating, newly mated queens face two options: independent colony foundation, in which a queen establishes a new colony alone, or dependent colony foundation, in which she joins an existing colony or founds a new colony with conspecifics. Independent foundation is highly risky; an estimated 99% of queens attempting it die (Cole 2009 and references therein), as the queen must find a suitable nesting site and then rear the first brood entirely on her own, surviving on fat reserves and catabolism of flight muscle accumulated during development. For this reason, only large queens are generally assumed to pursue this strategy, while small queens are expected to join established colonies, since the energetic demands of solitary founding would likely prove fatal.

Our data support this view: 75 of 78 newly mated queens collected during independent founding attempts were large MM-9a9a queens. We did observe two small P_2_P_2_–9r9r queens found alone under rocks, suggesting occasional independent founding attempts by microgynes; however, both died within days of collection, consistent with the idea that their reduced body size and limited fat reserves render this strategy largely inviable. Small heterozygous P_2_–9r queens face an additional disadvantage: a previous study suggests that such queens mating with M–9a or P_1_–9a males suffer severely reduced offspring fitness, with up to 75% of offspring dying due to epistatic interactions between supergenes on chromosomes 3 and 9 (Scarparo et al. 2026). Our data show that XP_2_–9a9r queens (where X can be any of the haplotype on chromosome 3) mate essentially at random across male haplotypes, and that 67% mate with 9a males (either M or P_1_ on chromosome 3). Combined with the small body size conferred by carrying at least one 9r haplotype, this mating pattern would condemn a substantial fraction of offspring to die, making independent colony foundation effectively impossible for this morph, and likely reinforcing dependence on established colonies.

We found no differences in the number of eggs laid by MM queens mated with males of different haplotypes, suggesting that the success of early colony foundation does not depend on mate genotype, consistent with findings in *F. selysi* (Avril et al. 2019b, Choppin et al. 2026). The fate of independently founded colonies headed by MM queens mated with P males remains unknown. In a previous study, P haplotypes were detected only sporadically across monogyne colonies (5 out of 60 colonies), with typically a single P-carrying individual per colony along with MM nestmates; notably, colonies analyzed by Scarparo et al. (2023) were mature and likely founded years prior. The combination of observed mating rates and the near-exclusive association of P haplotypes with polygyne colonies suggests that incipient colonies founded by MM queens mated with P males fail to persist or transition to polygyny, a trajectory supported by evidence from *F. selysi* (De Gasperin et al. 2025).

## Conclusions

This study reveals that the two supergenes in *Formica cinerea* do not act in isolation but jointly orchestrate a suite of traits (morphology, dispersal, mating strategy, and colony foundation) that together define distinct reproductive phenotypes (Figure 4). Each morph can be characterized by a coherent life-history profile that spans at least the first phases of the reproductive cycle until colony foundation. Monogyne macrogynes (MM–9a9a) are large, well-adapted for long-distance dispersal, mate assortatively, and can found colonies independently. Polygyne macrogynes (P_1_– 9a9a) appear intermediate, with reduced morphology suggesting shorter dispersal and potentially more flexible strategies than the monogyne counterpart, although the large body size could enable them to independently establish new colonies (Figure 4). In contrast, polygyne microgynes (P_2_–9r) are small, dispersal-limited, mate locally (sometimes within their natal nest), and cannot found colonies independently, relying instead on integration into existing colonies. Males retain broader mating flexibility, maintaining gene flow between morphs. At a broader level, our results highlight how genetic architectures can coordinate morphological, behavioral, and life-history traits into coherent strategies, thereby preventing the formation of disadvantageous combinations.

## Acknowledgements

We thank Marie Palanchon and Marco Molfini for their assistance in collecting *F. cinerea* queens, as well as Azariah Lopez and Elijah Muro for their help with genotyping. This work was supported by the US National Science Foundation Division of Environmental Biology grant #1754834 to AB and JP and grant #192252 to JP.

## Authors’ contributions

G.S: conceptualization, data curation, formal analysis, investigation, methodology, writing— original draft, writing—review and editing; A.B.: conceptualization, funding acquisition, investigation, methodology, project administration, writing—review and editing; J.P.: conceptualization, funding acquisition, investigation, methodology, project administration, supervision, writing—review and editing.

